# Serial femtosecond X-ray diffraction of HIV-1 Gag MA-IP6 microcrystals at ambient temperature

**DOI:** 10.1101/561100

**Authors:** Halil I Ciftci, Raymond G Sierra, Chun Hong Yoon, Zhen Su, Hiroshi Tateishi, Ryoko Koga, Koiwai Kotaro, Fumiaki Yumoto, Toshiya Senda, Mengling Liang, Soichi Wakatsuki, Masami Otsuka, Mikako Fujita, Hasan DeMirci

## Abstract

The Human immunodeficiency virus-1 (HIV-1) matrix (MA) domain is involved in the highly regulated assembly process of the virus particles that occur at the host cell’s plasma membrane. High-resolution structures of the MA domain determined using cryo X-ray crystallography have provided initial insights into the possible steps in the viral assembly process. However, these structural studies have relied on large and frozen crystals in order to reduce radiation damage caused by the intense X-rays. Here, we report the first XFEL study of the HIV-1 MA domain’s interaction with inositol hexaphosphate (IP6), a phospholipid headgroup mimic. We also describe the purification, characterization and microcrystallization of two MA crystal forms obtained in the presence of IP6. In addition, we describe the capabilities of serial femtosecond X-ray crystallography (SFX) using X-ray free-electron laser (XFEL) to elucidate the diffraction data of MA-IP6 complex microcrystals in liquid suspension at ambient temperature. Two different microcrystal forms of MA-IP6 complex both diffracted to beyond 3.5 Å resolution, demonstrating the feasibility of using SFX to study the complexes of MA domain of HIV-1 Gag polyprotein with IP6 at near-physiological temperatures. Further optimization of the experimental and data analysis procedures will lead to better understanding of the MA domain of HIV-1 Gag and IP6 interaction at high resolution and provide basis for optimization of the lead compounds for efficient inhibition of the Gag protein recruitment to the plasma membrane prior to virion formation.

## 1. Introduction

Soon after the first report of acquired immunodeficiency syndrome (AIDS) [1] its causative agent, human immunodeficiency virus (HIV) was isolated [2]. Since then the number of AIDS patients had increased worldwide and many patients had died due to the lack of effective treatment. Today we have a better understanding of how HIV replicates in a cell [3], and the life span of AIDS patients has been extended by antiretroviral therapy (ART) to that of normal people [3]. By the end of 2017, there were 36.9 million people living with HIV worldwide with 940,000 deaths from AIDS (https://www.who.int/hiv/en/). Some important discoveries related to HIV, so far, have come from crystallographic structure elucidation of HIV proteins (Figure 1A-D) [4]. Especially, first crystal structure of HIV protease transformed the treatment of HIV-1 patients [5–9]. These structural studies collectively led to understanding the function of HIV proteins at the molecular level and thus the development of anti-HIV drugs used in ART such as the first-generation HIV protease inhibitors[10]. Unfortunately, there is still no cure for HIV. However, there is significant advances in understanding the structure and function at a molecular level which bill bring the scientific community closer to a potential cure.

**Figure 1.**
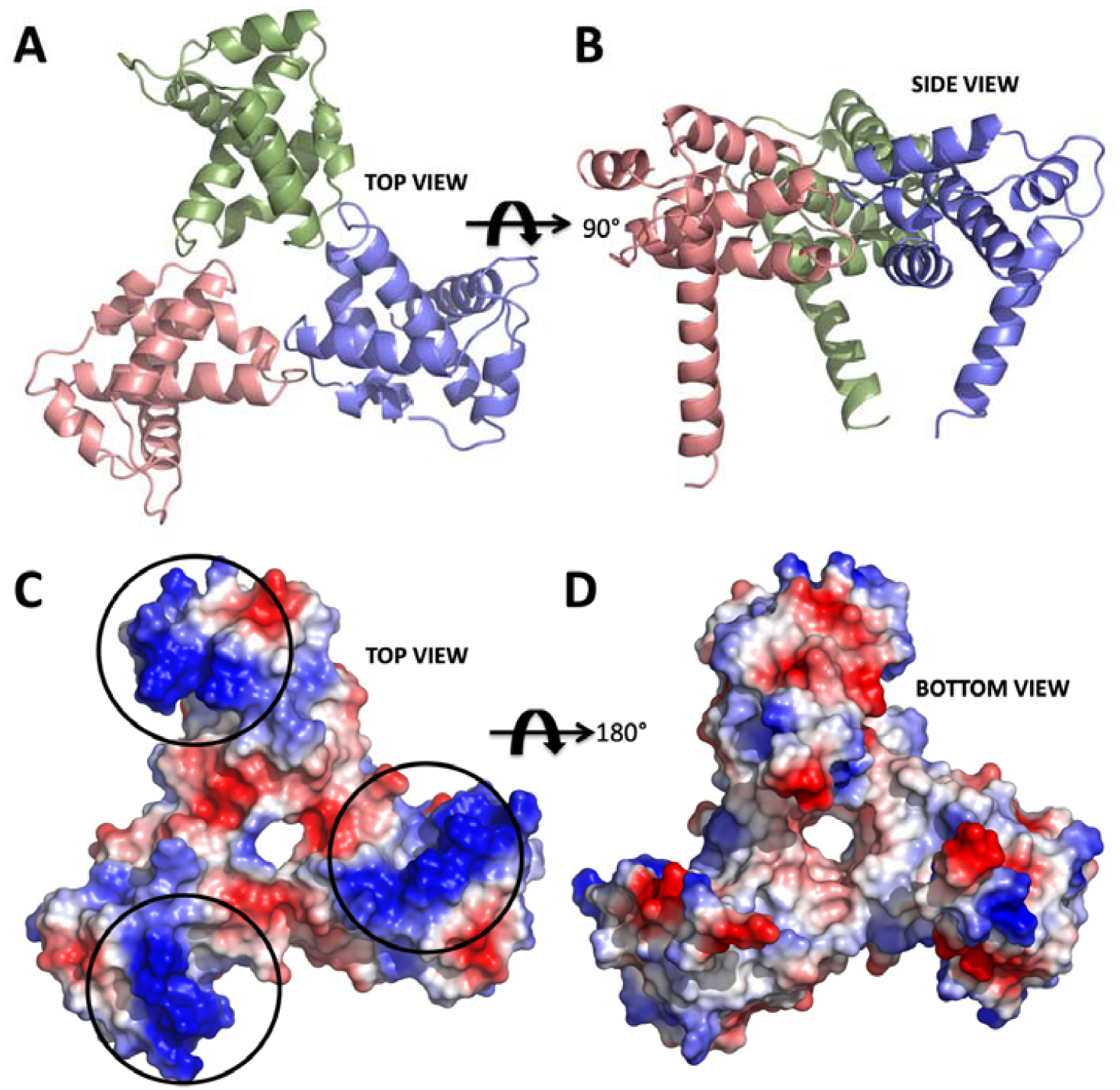
Structure of apo form HIV-1 matrix (MA) domain trimer at 2.3 Å resolution with R_free_: 0.322 and R_work_: 0.259 (adapted from Hill *et al*., 1996 PDB ID 1HIW). A) Top view of the MA trimer structure down the trifold symmetry axis with each subunit colored in salmon, green and slate. B) Side view of the MA trimer, same coloring scheme as in panel A rotated 90 degrees around the X-axis. C) Electrostatic surface potential of top part the MA trimer structure indicates the basic residues colored in blue are clustered on each of the three subunits. Black circles mark the putative binding sites for IP6. D) Electrostatic surface potential of the bottom side of the trimer indicating that IP6 only binds to the top part, due to the high electronegativity versus that of the bottom, which involves in membrane interaction via basic region.

Type 1 HIV (HIV-1), the major causative agent of the HIV pandemic, has only nine genes in its genome, including a structural gene *gag* that codes for Gag 55-kDa precursor (Pr55^gag^) protein. This protein is composed of four major domains, matrix (MA), capsid (CA), nucleocapsid (NC), and p6 successively from the N-terminus (Figure 2A). Pr55^gag^ plays the critical role in the virion release step. It is cleaved by a viral protease concurrently or immediately after virus budding, generating the four proteins, MA, CA, NC and p6. This cleavage is called maturation, in which the virus acquires infectivity, enables the virions to enter the target cells and eventually integrates reverse-transcribed viral DNA into the human genome [3]. Among the four domains, the most N-terminal MA domain mainly functions in membrane binding of Pr55^gag^ [11–14]and envelope (Env) incorporation into a virion [15]. The membrane binding of Pr55^gag^ is caused by the binding of the MA domain and D-*myo*-phosphatidylinositol 4,5-bisphosphate PI(4,5)P2, followed by insertion of the myristoyl moiety conjugated to the N-terminus of the MA domain into cellular membrane [13,16] (Figure 2A & B). Furthermore, cytoplasmic tail of Env gp41 interacts with the MA domain leading to the Env incorporation during the virion formation. Despite its importance throughout the replication cycle, the HIV-1 MA domain is not targeted yet by any of currently approved antiretroviral drugs [17–19]. Based on the structural information of the MA domain and PI(4,5)P2, we recently developed a non-natural derivative of PI(4,5)P2, named L-HIPPO, which binds to the MA domain in order to eradicate HIV [20,21]. Recent progress identifying cellular interactions of HIV-1 Gag has revealed that the MA domain of the Gag is capable of binding to inositol hexaphosphate (IP6) [20,22] (Figure 2B & C). Understanding the dynamic nature of the structural basis of these interactions at high-resolution may be achieved in the future and can provide new hypotheses for HIV therapy.

**Figure 2.**
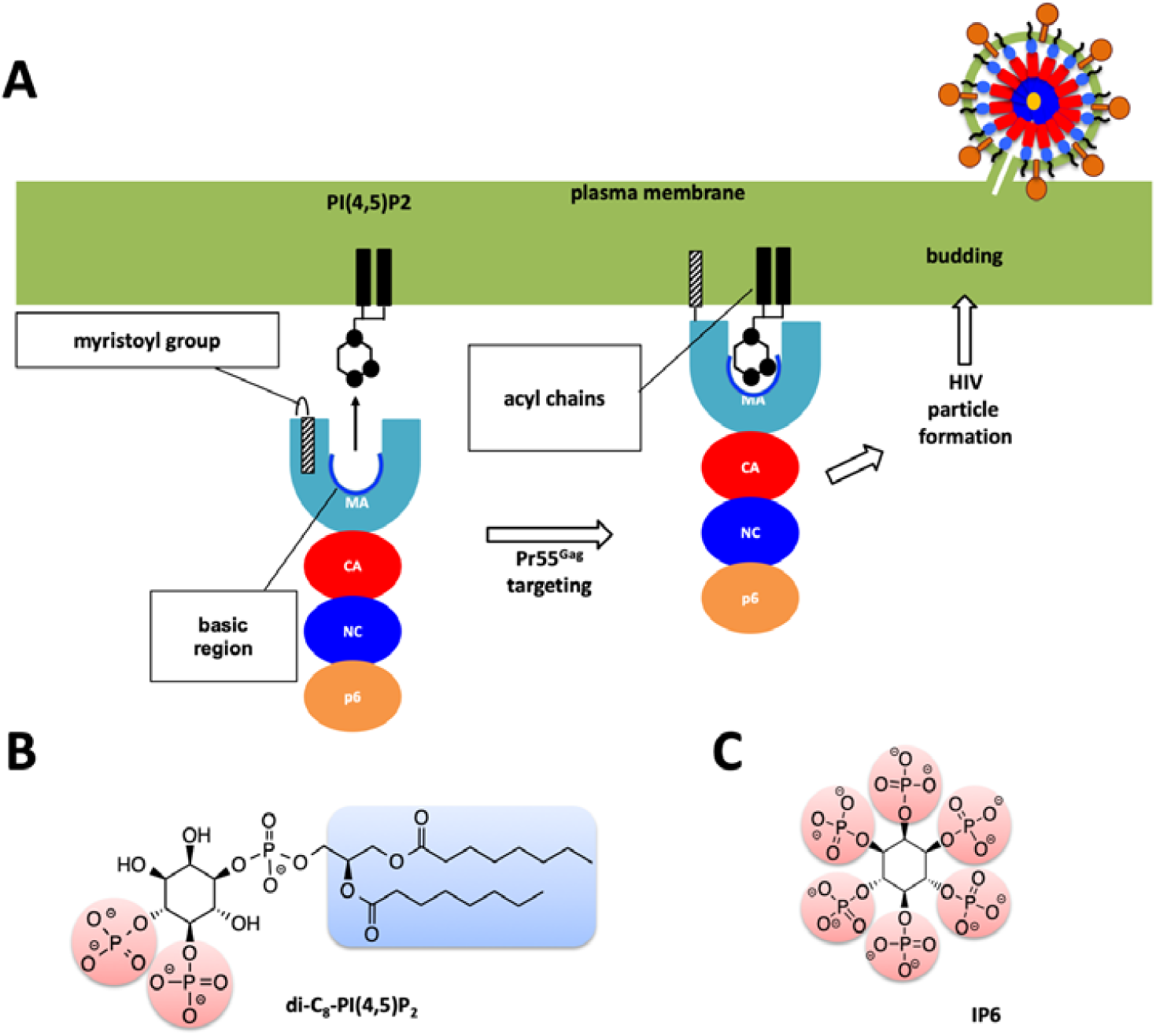
Role of MA domain in viral particle assembly and budding process. A) Pr55^Gag^ MA domain faces the host’s plasma membrane. For simplicity only one protomer of the trimeric Pr55^Gag^ complex has been shown in enlarged schematic view. Basic region of the MA domain which is marked with a blue curve which interacts with the acidic PI(4,5)P2 residue and facilitates the membrane attachment of the Pr55^Gag^ viral protein. Virion particle is not shown to scale and for simplicity Env proteins are not shown in the enlarged plasma membrane. B) Chemical structure of the PI(4,5)P2 molecule. C) Chemical structure of the IP6 molecule used in this work.

Structural studies of biological complexes in the near-physiological temperature range revealed previously-obscured conformations and provided a means to evaluate their local and large-scale dynamics [23–25]. Serial Femtosecond X-ray Crystallography (SFX) is a new technique that uses X-ray Free-Electron Lasers (XFELs) to determine protein structures at ambient or cryogenic temperature. XFEL lightsources are capable of generating pulses of X-rays spanning tens of femtoseconds in duration and exceeding the brightness of current synchrotrons [26,27]. The linac coherent light source (LCLS) at SLAC national accelerator laboratory was the first such XFEL capable of producing X-ray pulses of 10^12^ photons at photon energies ranging from 500 eV to 12.7 keV with a duration of several to a few hundred femtoseconds, which is about 100 million to a billion times brighter than the synchrotron X-rays [28–30]. SFX harnesses these pulses to probe microcrystals at ambient temperature and quickly emerged as a promising new method to complement synchrotron-based crystallography studies [26,27,31–33]. The most common SFX approach is to deliver sub-micron to 20 micron size crystals flowing in a liquid suspension to the interaction point, at which the extremely short and brilliant X-ray pulses interact with the microcrystals and produce diffraction patterns before Coulomb explosion of these microcrystals [34–37]. Matching crystal size to the beam size typically minimize the background and maximizes the signal quality. The ability of the ‘diffract-before-destroy’ approach to obtain high-resolution data was first demonstrated by the 1.9 Å resolution structure of lysozyme and the 2.1 Å resolution structure of cathepsin B [38–40]. The potential of this approach for the study of large macromolecular complexes has also shown great promise (see for example [27,41,42]).

Until recently, the X-ray crystallographic studies of HIV proteins were limited to synchrotron cryo X-ray crystallography. One particular important development is the recent advances in micro Electron Diffraction (microED) technique which can use submicron crystals to produce high-resolution structures [43]. There are not many examples of XFEL structures of HIV proteins except one recent report on HIV-1 envelope (Env) [44]. One of the biggest challenges to perform SFX studies is to produce a sufficient amount of high-quality protein microcrystals that allows for the determination of high-resolution structures. In SFX, diffraction data are collected from microcrystals within their native crystallizing media using extremely bright, extremely short and highly coherent XFEL pulses (Figure 3). The short duration of X-ray pulses, on the order of tens of femtoseconds, produces high resolution diffraction patterns before the onset of radiation damage of the crystals. The ability of the diffraction-before-destruction principle represents the basis of SFX that enables to collect high-resolution data at room temperature [38]. SFX also allows collection of time-resolved diffraction patterns during of the reactions that are linked to enzymatic reactions (the bond making/breaking steps and structural conformational changes) occur on the timescale from femtoseconds to milliseconds. Taken together, SFX can enable a better understanding of reaction or binding intermediates in previously unobserved detail.

**Figure 3.**
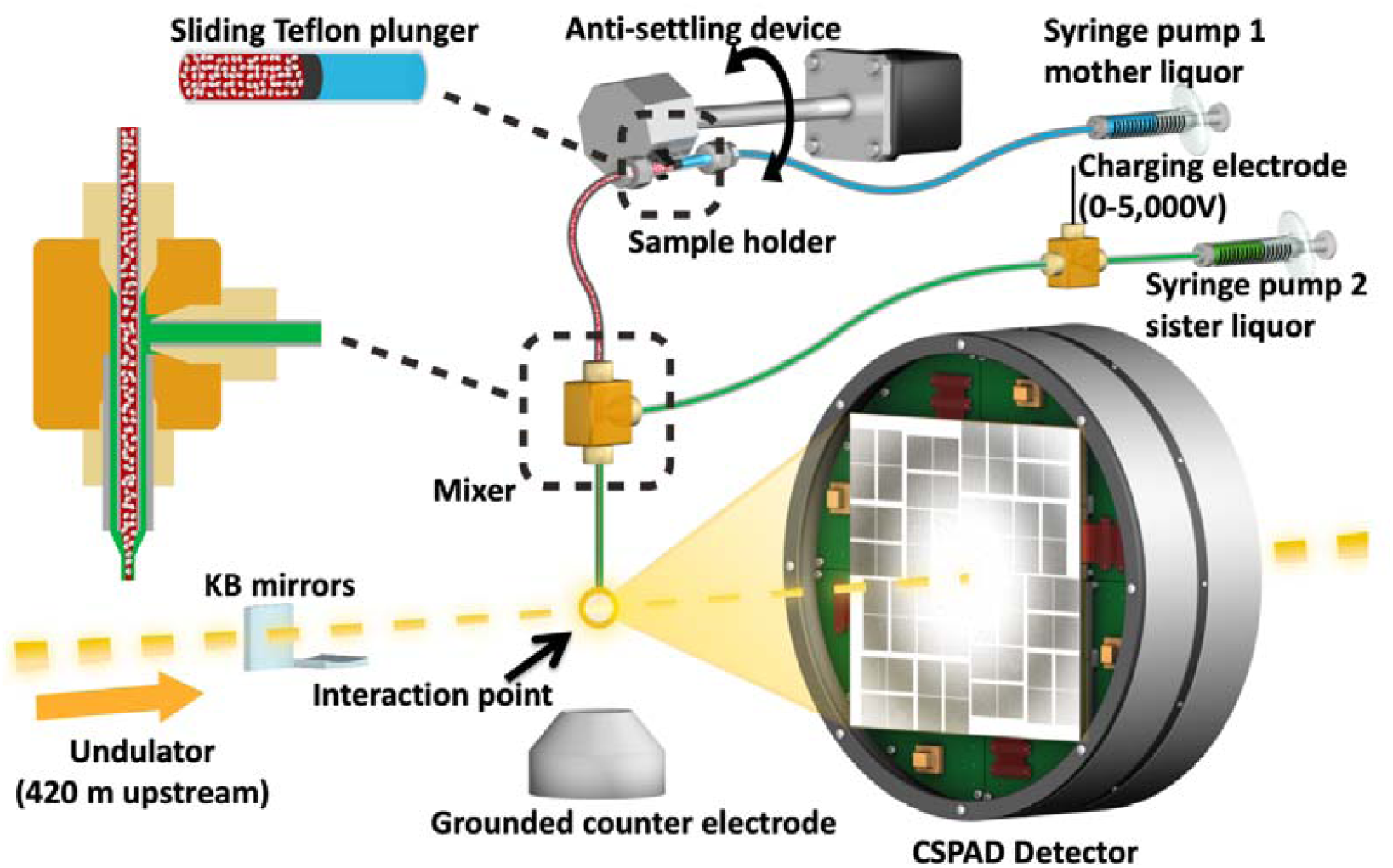
Image of the coMESH injector setup at the CXI instrument of the LCLS. The liquid jet, comprising MA-IP6 microcrystals and their mother liquor (20% w/v PEG3350 and 100mM MES pH 6.5; colored in red), flowed in the continuous inner capillary (100 μm × 160 μm × 1.5 m; colored in gray). The sister liquor containing mother liquor supplemented with 20% MPD (colored in green) was charged by a high voltage power supply (0–5,000 V) for electro-focusing of the liquid jet. A microfluidic tee junction (indicated within the dashed orange square) joined the two capillaries (colored in gray) concentrically. The sample reservoir had a Teflon plunger (colored in black) which separated the sample reservoir from the driving fluid (water, colored in light blue). The reservoir was mounted on an anti-settling device which rotated, at an angle, about the capillary axis to keep the protein crystals suspended homogenously in the slurry. The liquid jet and the LCLS pulses with 1×1 μm^2^ focus interacted at the point indicated by the orange circle.

Here we present the feasibility of such structural studies on HIV-1 interaction with phospholipid membrane budding using XFELs. We also describe the experimental procedures from purification and characterization of HIV-1 MA protein, its large-scale co-crystallization with IP6 in two crystal forms, and efficient delivery of these crystals for ambient-temperature diffraction data collection through an SFX experiment. Using 40-femtosecond pulses at 9.5 keV at the Coherent X-ray Imaging (CXI) instrument at LCLS, we obtained diffraction from MA-IP6 microcrystals prior to the onset of radiation damage induced by the X-rays [45]. The microcrystals were introduced to the X-ray beam in a liquid suspension with a concentric electrokinetic microfluidic sample holder (coMESH) injector [23].

## 2. Results

For the SFX experiments, the hanging drop crystallization conditions were optimized to favor the formation of microcrystals by increasing the number of drops to 15-20 per coverslide. After harvesting in the same mother liquor, microcrystals were pooled, and suspensions were pre-filtered through a 40 μm nylon mesh filter to remove large particles and aggregates (Figure 4). Final sample slurry contained crystalline mixture of 1×1×5 μm^3^ to 5×5×15 μm^3^ size range which is measured by light microscopy. Crystals were kept at 293 K before being introduced into the LCLS beam in a thin liquid jet using the coMESH injector (Figure 3) at flow rates between 1-3 μl/min. Total of 500 μl sample used for each dataset. The sheath liquid contained the mother liquor, but added a 20% v/v of MPD, and was charged between 3-5 kV and flowrates varied between 1-3 μl/min in order to maintain jet stability and maximize hit rate while monitoring with *OnDa* [46]. The 40 femtosecond X-rays pulses intercepted the continuous jet at 120 Hz, with a pulse energy between 2-3 mJ at a wavelength of 9.5 keV.

**Figure 4.**
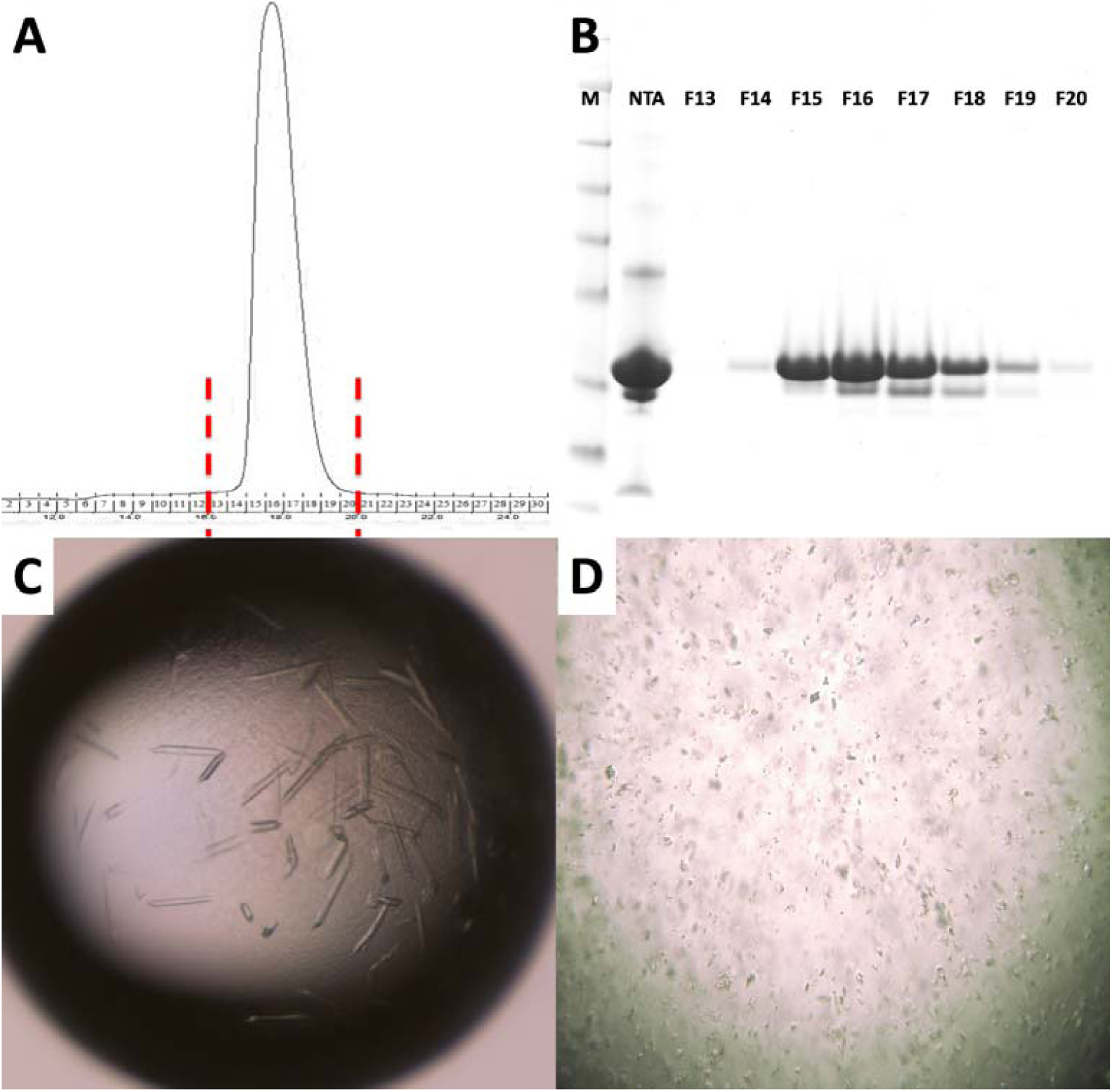
Purification and characterization of MA domain of A) Size exclusion chromatography of apo MA domain yields a monodisperse pattern indicating it is stable as a monomer. B) 15% SDS PAGE gel shows the final purity level of the of the MA domain that was used for crystallization experiments. M: marker, NTA: Elute from Ni-NTA column, F13-F20 are the fractions 13-20 from S200 column. C) Image of HIV-1 MA - IP6 co-crystals before and D) after filtration through 40-micron Millipore nylon mesh filter.

Diffraction data were recorded using a Cornell–LCLS pixel array detector (CSPAD) detector [47]. 180,612 (12%) potential crystal hits were identified from 1,530,202 diffraction patterns using a crystallography software called *Psocake* [48,49]. Diffraction was observed to a resolution beyond 3.5 Å (Figure 5).

**Figure 5.**
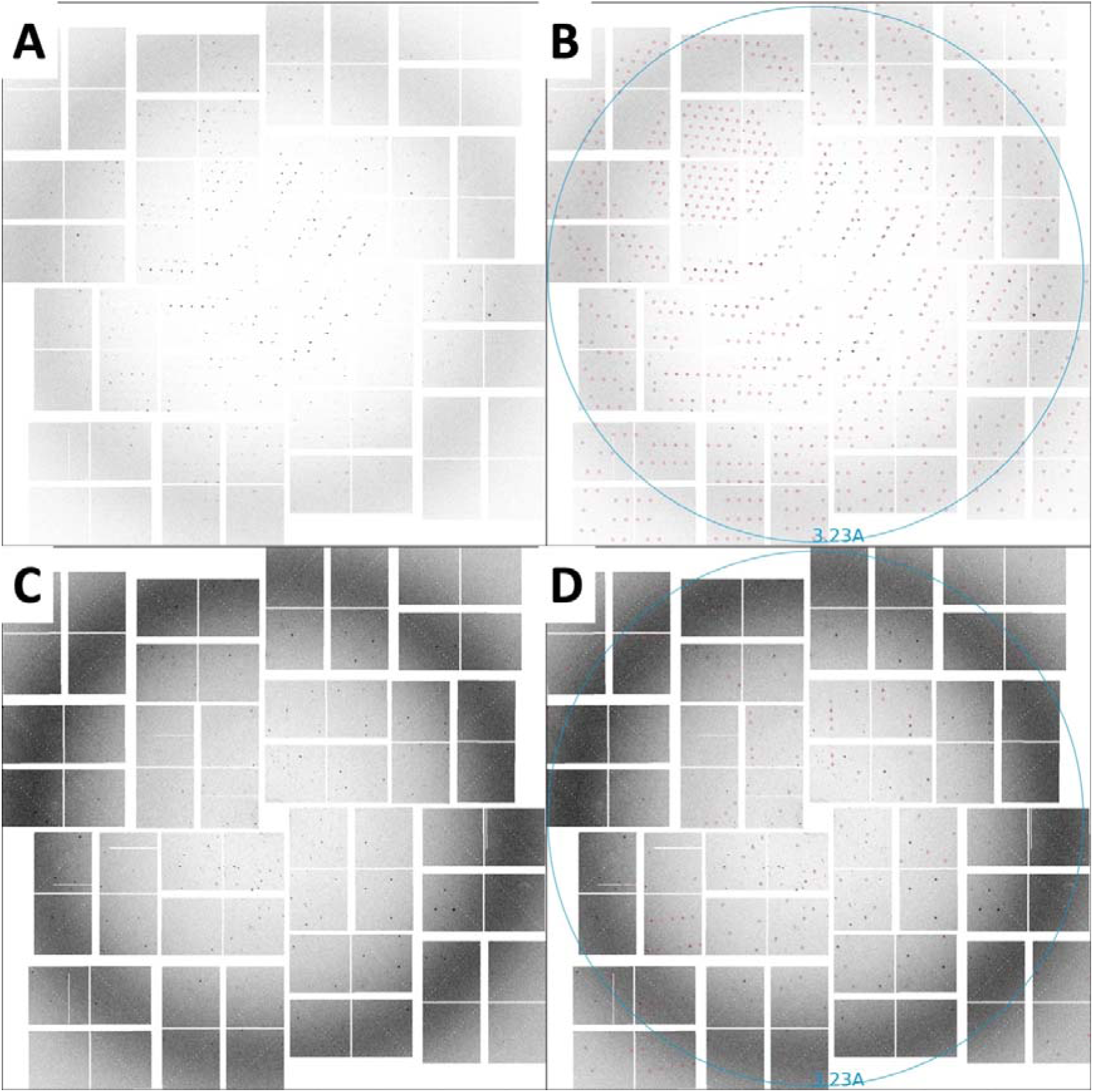
SFX diffraction image collected on a CSPAD detector. A) Diffraction spots from rod-shaped crystals extending to beyond 3.5 Å resolution, with unit-cell parameters *a* = *b* = 96.5, *c* = 91.1 Å, α = β = 90°, γ = 120°. B) The same image as in panel A with the reflection predictions after indexing circled in red. C) Diffraction spots from triangular-shaped crystals extending to beyond 3.5 Å resolution, with unit-cell parameters *a* = *b* = 96.8, *c* = 91.8 Å, α = β = 90°, γ = 120°. D) The same image as in panel C with the reflection predictions after indexing circled in red.

## 3. Discussion

The observed lower-than-expected resolution of HIV-1 MA - IP6 complex may be accounted for, in part, by the treatment of microcrystals in the particular experimental set up available at the time of the experiment. Relatively large size crystals up to 5×5×15 μm^3^ compared to 1×1 μm^2^ beam size and high background due to low dynamic range of CSPAD detector could potentially increase the background while lowering the signal quality. Potentially many factors can affect the data quality of an SFX experiment. A large unit cell size and limited number of crystal contact points with high solvent content can lead to a more physically delicate crystal, which can be damaged when transferred into reservoirs or passed through filters [42]. Samples can also suffer incompatibility with the injection method and the sample environment. These potential shortcomings were not screened for in this experiment which simply served as a proof-of concept. From prior experiences, the authors suspect that growing these large crystals and harvesting them from plates to reservoirs, can clearly show visible crystal degradation such as jagged edges, clumps, bent crystals etc. leading to poor diffraction data. Indeed, crystals had been subjected to continuous pipetting to remove them from the glass cover slides which caused mechanical shearing forces before injection across the LCLS beam. These repeated physical contacts with crystals can damage the packing arrangement of the delicate MA-IP6 complexes in the crystal lattice, introduce additional mosaicity to the crystals and thereby lower the resolution limit. During beamtime large numbers of i.e. 54,178 diffraction patterns for MA-IP6 rod and 126,434 diffraction patterns for MA-IP6 triangle crystals had been recorded (Table 1). To obtain the final sample slurry, hundreds of microliter size drops has been combined and this pooling process most likely caused heterogeneity and non-isomorphism in unit cell parameters. Despite our exhaustive efforts non-isomorphic unit cell parameters so far precluded the successful merging of the data to recover electron density. Future experiments will employ a gentler batch method in combination with density gradient separation of microcrystals by size rather than filtering, to improve resolution and lower the non-isomorphism to streamline the downstream dataprocessing [23]. Crystallization protocols can also be optimized, including the examination of different crystal forms and geometries to determine the optimum shape and size of the microcrystals for future SFX studies, thereby eliminating the need for filtering.

**Table 1.** *Psocake* hit finding parameters.

| Peak finding algorithm | Peak criteria | Min. peaks |
| --- | --- | --- |
| Adaptive | Amax_thr 300, Atot_thr 600, Peak size 2-30, Son_min 10, Rank 3, Radius 3, Dr 2 | 15 |

Most of the temperature dependent dynamics such as rotation and conformational flexibility of the side chains of the proteins are captured in frozen conformations at cryogenic temperatures. This impedes the understanding of binding dynamics of the HIV-1 MA-IP6 complex such as order of formation of H-bonds and coupled structural conformational changes will remain unclear. Furthermore, the requirement of synchrotron X-ray crystallography for large crystals hampers the structure of HIV-1 MA - IP6 complex in the different intermediate states of the binding process that might limit the crystal growth. Often, larger crystals need to be frozen at very low temperatures that also increase mosaicity and lower resolution with increasing radiation damage that negatively impact the quality of diffraction data. However, recent upgrades in microfocus synchrotron beamlines optics and direct photon count pixel detectors allow data collection from as small as 1-micron size crystals, however, this method still necessitates the cryocooling. The dynamic structure of the HIV-1 Gag MA domain and its complexes with IP6 would have benefit significant advantage from the ability for structural studies at temperatures closer to the physiological condition in which these processes take place.

## 4. Materials and Methods

### 4.1 Cloning and overexpression of the HIV-1 Gag MA domain

His10-tagged HIV-1 MA gene was cloned into pRSF-1b vector and grown in Luria-Bertani (LB) media in the presence of 50 μg/ml kanamycin antibiotic for 4 hours. Expression of the MA domain induced by 0.5 mM isopropyl β-D-1-thiogalactopyranoside (IPTG), incubated overnight in *E. coli* BL21 (DE3) cells at 16° C. Cells were harvested in lysis buffer containing 20 mM 2-Amino-2- (hydroxymethyl) propane-1,3-diol (Tris)-HCl pH 8.0, 0.5 M NaCl, 30 mM imidazole, 5 mM β-mercaptoethanol and lysed by sonication. Cell debris and membranes were pelleted at 18000 rpm by centrifugation. The remaining supernatants, which contain the soluble MA fraction, were pooled and loaded on to Nickel-Nitriloacetic acid (Ni-NTA) column and washed with 10X column volume of wash buffer containing 20 mM Tris-HCl pH 8.0, 1 M NaCl, 1 M (NH_4_)_2_SO_4_, 30 mM imidazole and 5 mM β-mercaptoethanol. Bound MA fractions were eluted with elution buffer containing 20 mM Tris-HCl pH 8.0, 0.5 M NaCl, 300 mM imidazole and 5mM β-mercaptoethanol, and then buffer exchanged and the His_10_-tag was cleaved off with Tobacco Etch Virus (TEV) protease at pH 8.0. The His-tagged TEV protease and uncleaved His_10_-tagged MA was removed by the Ni-NTA column. Two ml of the fraction was mixed with a denaturing buffer (20 mM Tris-HCl, pH 8.0, 1 M NaCl, 1 M (NH_4_)_2_SO_4_, 1 mM dithiothreitol (DTT), 6 M urea) and dialyzed in the buffer to remove contaminating nucleic acids for overnight. After concentrating the dialyzed sample to 1 ml, polypeptides were refolded through a size exclusion column Superdex200 10/300 increase (GE Healthcare) in the buffer (20 mM Tris-HCl pH 8.0, 150 mM NaCl, 1 mM DTT). The fractions containing MA domain were confirmed by SDS-PAGE and concentrated. The sample was prepared at the High Energy Accelerator Research Organization, Tsukuba, Ibaraki, Japan and delivered to the LCLS, Menlo Park, CA, USA for micro-crystallization and data collection (Figure 4A &B).

### 4.2 Crystallization of the HIV-1 Gag MA domain

Purified HIV-1 Gag MA proteins were then used in co-crystallization with IP6 at room temperature by the hanging-drop method using a crystallization buffer containing 20% polyethylene glycol 3350 (PEG 3350) as precipitant and 100 mM MES-NaOH pH 6.5. Microcrystals were harvested in the same mother-liquor composition, pooled to a total volume of 3 ml (a representative hanging drop is seen in Figure 4C) and filtered through 40-micron Millipore mesh filter (Figure 4D). The concentration of crystal was around 10^10^-10^11^ per milliliter viewed by light microscopy.

### 4.3 XFEL X-ray delivery and detector

An average of 2.64 mJ was delivered in each 40-fs pulse contained approximately 10^12^ photons with 9.51 keV photon energy with 1×1 μm^2^ focus of X-rays. Single-pulse diffraction patterns from HIV-1 MA-IP6 microcrystals were recorded at 120 Hz on a CSPAD [47] detector positioned at a distance of 217 mm from the interaction region.

### 4.4 Injection of HIV-1 MA-IP6 microcrystals into an XFEL and diffraction data collection

A crystalline slurry of HIV-1 MA-IP6 microcrystals kept at room temperature flowing at 2 μl/min was injected into the interaction region inside the front vacuum chamber at the LCLS CXI instrument using the coMESH injector (Figure 3). Due to presence of large crystals in the MA-IP6 samples, prior to the experiment, the coMESH injector required filtered sample before injection through a 100-micron inner diameter capillary size to prevent clogging.

### 4.5 Hit Finding

The SFX diffraction data collected from two different crystal forms (rod and triangle shapes) at LCLS were processed using *Psocake* software [48,49], yielding two complete datasets. A diffraction pattern was deemed a hit if at least 15 peaks were found. A total of 54,178 diffraction patterns for MA-IP6 rod and 126,434 diffraction patterns for MA-IP6 triangle crystals were recorded as crystal hits (Table 1). The peak finding parameters for the chamber are also summarized in Table 1.

### 4.6 Indexing

*CrystFEL’s indexamajig* program [50] was used to index the crystal hits. Two rounds of indexing were performed on each of the datasets. Initial indexing results indicated that that the space group was most likely hexagonal P6 with a = 96.55 Å, b = 96.78 Å, c = 91.02Å and α = β = 90°, γ = 120°. Given the target unit cell, the indexing results were accepted if the unit cell lengths and angles were within 5% and 1.5 °, respectively (Table 2). The final iteration yielded 25,501 (47%) and 56,861 (45%) indexed patterns for MA-IP6 ROD and MA-IP6 TRIANGLE, respectively. Representative patterns are shown in Figure 5. Figures of merit for merged intensities CC_1/2_ and R_split_ for MA-IP6 Rod and MA-IP6 Triangle are shown in Figure 6. The merged intensities were symmetrized with point group 6/mmm and the estimated resolution is around 3.3 Å.

**Table 2.** *CrystFEL* indexing parameters.

| Indexing algorithm | Integration radii | Unit cell tolerance |
| --- | --- | --- |
| Mosflm, dirax | 3,4,5 | Axes lengths=5%<br>Axes angles=$1.5^\circ$ |

**Figure 6.**
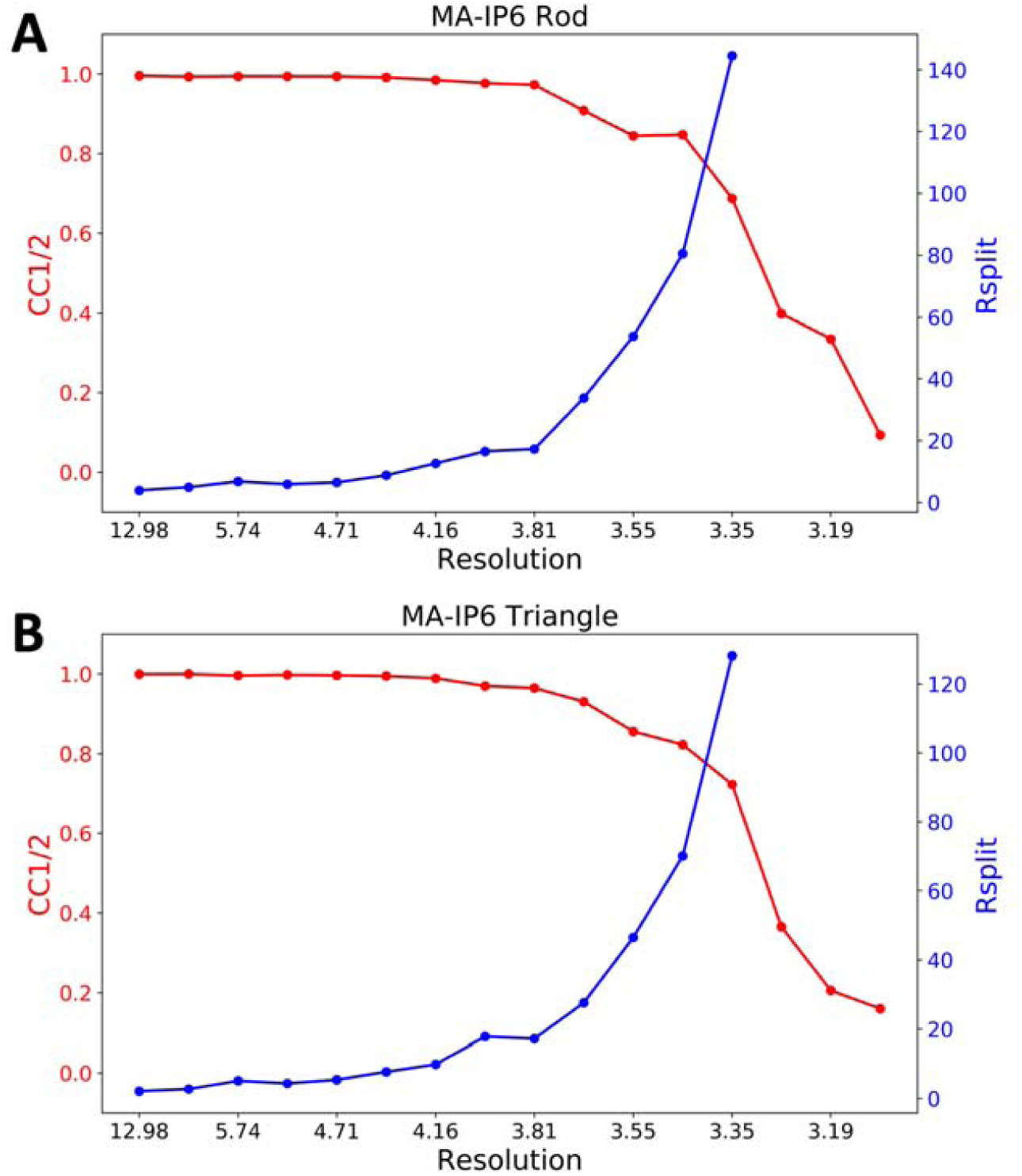
Figures of merit plot: CC_1/2_ (red) and R_split_ (blue) versus resolution for A) MA-IP6 Rod and B) MA-IP6 Triangle.

## 5. Concluding remarks

Diffraction patterns of MA-IP6 microcrystals were recorded beyond 3.5 Å resolution. It was possible to determine the unit-cell parameters using *Psocake* [48,49]and the *CrystFEL* [51,52]software suites. The unit-cell parameters could be estimated (Figure 5). These results demonstrate the feasibility of conducting HIV-1 MA-IP6 complex structural studies using XFELs, which hold great promise for a more comprehensive understanding of MA and IP6 interaction by performing time-resolved studies unique to XFELs, in which structural intermediates may be imaged. Recent SFX studies of an adenine riboswitch aptamer domain mixed with its substrate immediately prior to probing captured dynamics of the reaction at ambient temperature and revealed conformational changes that also induced a conversion of the space group *in crystallo* [53]. Such findings indicate that mix-and-probe time-resolved SFX can offer opportunities to probe macromolecular complexes using microcrystals, either as static structures or as they undergo biologically relevant reactions.

## Author Contributions

H.D., H.I.C., and M.F. designed and coordinated the project. H.I.C., H.T., K.K., and F.Y. established the protein expression and purification method for the crystallization. T.S. supervised the crystallographic experiment. H.D. and H.I.C prepared and characterized the samples. H.D., C.H.Y. and Z.S. analyzed data. R.G.S., H.D. and H.I.C. built the co-MESH injector, helped with data collection. R.G.S., and M.L. prepared the beamline and ran the CXI instrument. H.D., M.F., H.I.C., and M.O. wrote the manuscript with input from all of the authors.

## Acknowledgments

This work was supported in part by a Grant-in-Aid for Scientific Research (C) from the Japanese Society for the Promotion of Science (JSPS) (17K08861), Grant-in and Aid for JSPS Research Fellow (17J11657), and the Platform for Drug Discovery, Informatics, and Structural Life Science (Ministry of Education, Culture, Sports, Science and Technology (MEXT), Japan Agency for Medical Research and Development (AMED)) and Basis for Supporting Innovative Drug Discovery and Life Science Research from Japan Agency for Medical Research and Development (AMED). Portions of this research were carried out at the Linac Coherent Light Source (LCLS) and Stanford Synchrotron Radiation Lightsource (SSRL) at the SLAC National Accelerator Laboratory. Use of the LCLS and SSRL is supported by the U.S. Department of Energy (DOE), Office of Science, Office of Basic Energy Sciences (OBES) under Contract No. DE-AC02-76SF00515. The LCLS is acknowledged for beam time access under experiment no. cxils9717. HD acknowledges support from NSF Science and Technology Center grant NSF-1231306 (Biology with X-ray Lasers, BioXFEL). Parts of the sample injector used at LCLS for this research was funded by the National Institutes of Health, P41GM103393, formerly P41RR001209. H.D. acknowledges valuable discussions with Aiko Takeuchi, Kenji Dursuncan, Emi Satunaz and Michelle Young. We thank Gregory Stewart of SLAC National Accelerator Laboratory for excellent technical assistance with creating graphics for Figure 3.

